# Multigenerational Tracking of Reef-Building Corals Using CRISPR-Cas9 induced Genetic Barcodes and eDNA Metabarcoding

**DOI:** 10.1101/2025.09.11.675722

**Authors:** Max Moonier, Line Bay, Yui Sato, Véronique Mocellin, Phillip Cleves, Ryan Lister, Luke Thomas

## Abstract

Widespread biodiversity loss driven by human activity has intensified global efforts to restore degraded ecosystems. Yet a key challenge remains: how to track restored individuals and their offspring over time and space to assess the true impact of restoration? This is especially pressing for coral reefs, which support a quarter of all marine species, are in severe decline globally, and are now the focus of growing restoration initiatives worldwide. Here, we demonstrate a novel approach that combines genome editing and environmental DNA (eDNA) monitoring to enable scalable, non-invasive tracking of individual corals and their offspring. Using CRISPR-Cas9, we introduced unique genetic barcodes into non-coding regions of the *Acropora millepora* genome. These barcodes - stable, heritable, and distributed across the genome - will allow for individual-level identification over multiple generations. We show that these barcodes can be reliably detected in surrounding seawater using eDNA metabarcoding, offering a powerful, non-destructive tool for tracking corals *in situ*. Together, this approach provides a proof-of-concept for precision monitoring of restoration outcomes, with broad applicability across species and ecosystems.

## Introduction

Global biodiversity is declining due to direct anthropogenic pressures such as habitat destruction, pollution, and overharvesting, as well as the indirect effects of climate change^1–3^. In response, conservation strategies have shifted from passive protection to active restoration. In marine systems, this includes releasing hatchery-reared juveniles to rebuild fisheries (e.g. salmon, lobster, abalone) and reintroducing habitat-forming species like mangroves, kelp, and corals to support ecosystem services^4–9^. On land, similar efforts include rewilding, translocations, and large-scale planting of foundational species, such as trees and prairie grasses, to restore ecological function and biodiversity^10–12^.

Monitoring the long-term success of interventions is crucial to understanding what practices are working and where. However, measuring the success of interventions requires tools to reliably distinguish restored individuals from wild conspecifics, enabling tracking of their survival, reproduction, and dispersal over time, which has remained a challenge especially in species with long lifespans and highly dispersive offspring ^13,14^. Traditional approaches to access survival in the marine environment rely on physical tags monitored with diver-based surveys, which are logistically challenging, costly, and often limited in their ability to track individuals over ecologically relevant time and space^15–18^. Genetic methods like Parentage-Based Tagging and Genetic Stock Identification offer alternatives^19,20^, but require extensive multi-locus genotyping, large reference datasets, and repeated sampling of broodstock, limiting their scalability^21,22^. Furthermore, these tools are limited in their ability to track restored individuals and their offspring across multiple generations - a critical limitation for evaluating long-term restoration outcomes, particularly in species with sessile adults and highly dispersive offspring^23^.

Coral reefs are declining at an alarming rate due to a variety of anthropogenic stressors, despite their ecological and economic importance^24–30^. In response, restoration strategies are increasingly focused on selectively propagating large quantities of coral, enabled by advances in aquaculture, breeding, and molecular genetics^31–38^. However, tracking restored individuals and their offspring across space and time remains a key challenge^39,40^, especially given corals’ long lifespans (hundreds to thousands of years in many species^41,42^) and widespread offspring dispersal^43^.

Here, we present a proof-of-concept for combining CRISPR-Cas9 genome editing^44–46^ and environmental DNA (eDNA) metabarcoding to create and detect unique, heritable genetic barcodes in the ubiquitous reef-building coral, *Acropora millepora* (Fig. 1A). By targeting repetitive non-coding genomic regions, we generated individuals with diverse and persistent barcode combinations that were detectable in surrounding seawater. This approach lays the groundwork for scalable, non-invasive monitoring of restored individuals and their progeny over ecologically relevant timescales.

**Fig. 1:**
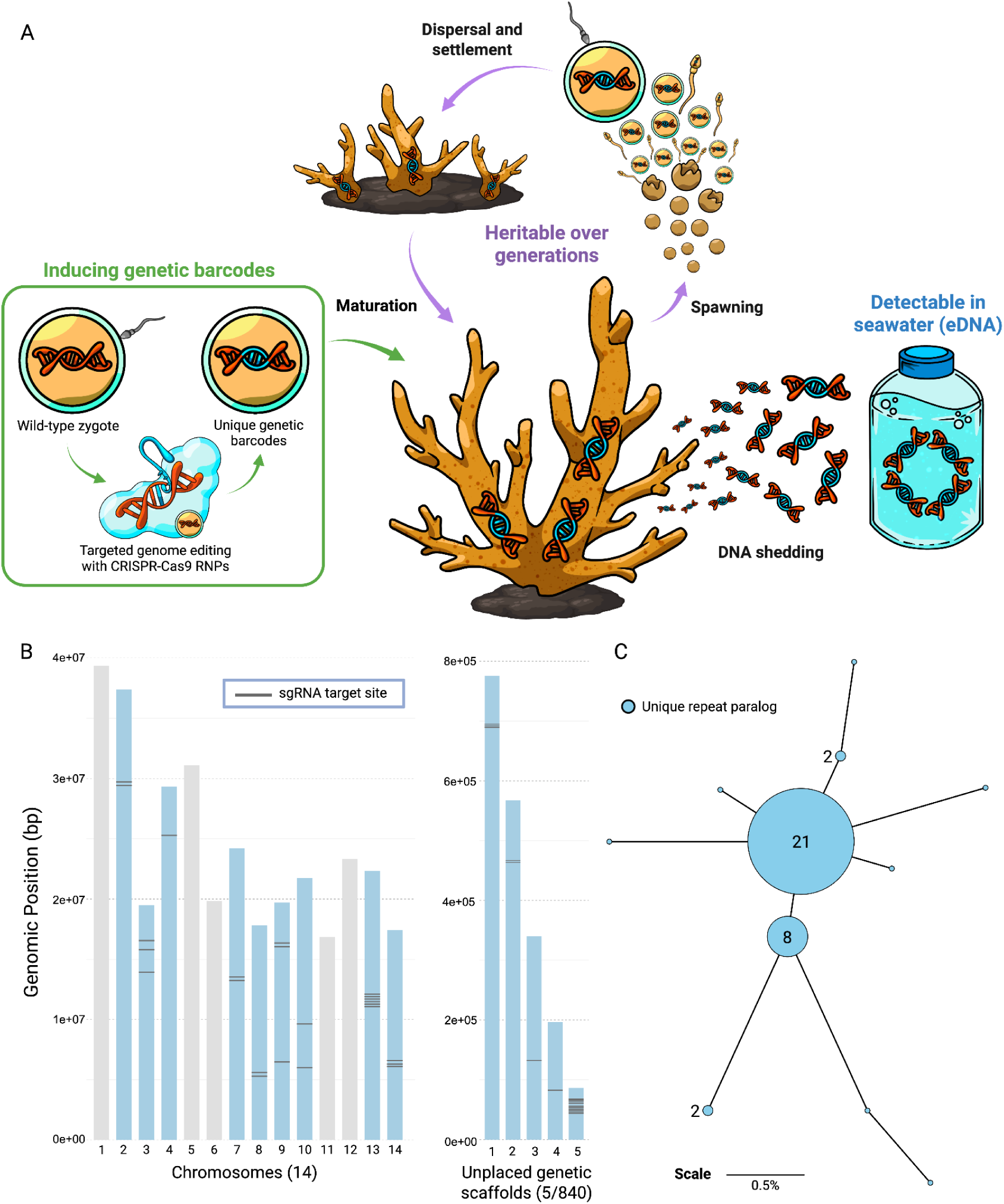
Distribution and sequence diversity of 41 repetitive target loci. (A) Schematic summary of CRISPR-Cas9 mediated genetic barcode induction and application. Wild-type (WT) repetitive target loci are represented by red DNA and unique genetic barcodes are represented as the blue section of DNA inserted into the WT repetitive target. (B) Map of all 14 chromosomes and 5/840 unplaced genetic scaffolds in the *A. millepora* reference genome. Chromosomes/scaffolds containing sgRNA target sites are coloured in blue. The genomic location of the 41 sgRNA target sites are represented with horizontal grey bars. (C) Haplotype network showing sequence difference between 11 unique target paralogs. Branch lengths are proportional to percent sequence difference between paralogs. Paralog abundance is indicated for paralogs with abundance >1 and node sizes are scaled accordingly.

## Results

### Identifying a repetitive target sequence

To increase the frequency and detectability of our CRISPR-Cas9-induced genetic barcodes, we targeted a repetitive DNA region spread across the genome, which is noncoding and putatively non-functional. We designed a set of PCR primers that amplified a 234 bp region spanning the repeat that contained target sites for three CRISPR-Cas9 single guide RNAs (sgRNA) (Fig. S1). We then used the basic local alignment search tool (BLAST) to query the 234 bp amplicon sequence against the *A. millepora* reference genome on NCBI (accession number PRJNA633778), which identified 41 loci in the reference genome that could be amplified by our PCR primer set, all of which had a 100% sequence match for at least 2 of the 3 sgRNAs and none of which overlapped any coding sequences or annotated regions of the reference genome. We mapped these 41 target sites to the reference genome and found this repeat to be wide-spread and present on 9 of the 14 *A. millepora* chromosomes along with 5 unplaced scaffolds (Fig. 1B). Sequence analysis revealed that the 41 repeat loci are composed of 11 unique paralogs harboring small sequence differences (Fig. S2), though they are dominated by two common haplotypes (Fig. 1C).

### Efficient CRISPR-Cas9 editing of the target repeat in *A. millepora*

To generate unique genetic barcodes, we injected *A. millepora* zygotes with a mix containing preassembled sgRNA/Cas9 ribonucleoprotein targeting the repeat and a fluorescent tracer. Control larvae were injected with the same mix but lacking the sgRNAs. We sequenced a subset of treated larvae (N=8), all of which exhibited substantial insertions and deletions (indels) within the CRISPR editing window (Fig. S3), with sgRNA 2 exhibiting substantially higher editing efficiency than sgRNAs 1 and 3 (Fig. 2A). We detected indels (in relation to the most dominant paralog sequence) in 47% of recovered reads in treated larvae, compared to 1% in control (Fig. 2B). We then clustered unique sequences into amplicon sequence variants (ASVs) and compared ASVs found in larvae to those found in parents. While control larvae contained only the WT ASVs, treatment larvae had up to 70 unique ASVs (mean ± SE: 41.7 ± 14.2 unique ASVs) (Fig. 2C) representing the unique inserted barcodes. Together, these results indicate high CRISPR-Cas9 editing efficiency at our repetitive target, generating a high proportion of barcoded sequences originating from numerous independent editing events.

**Fig. 2:**
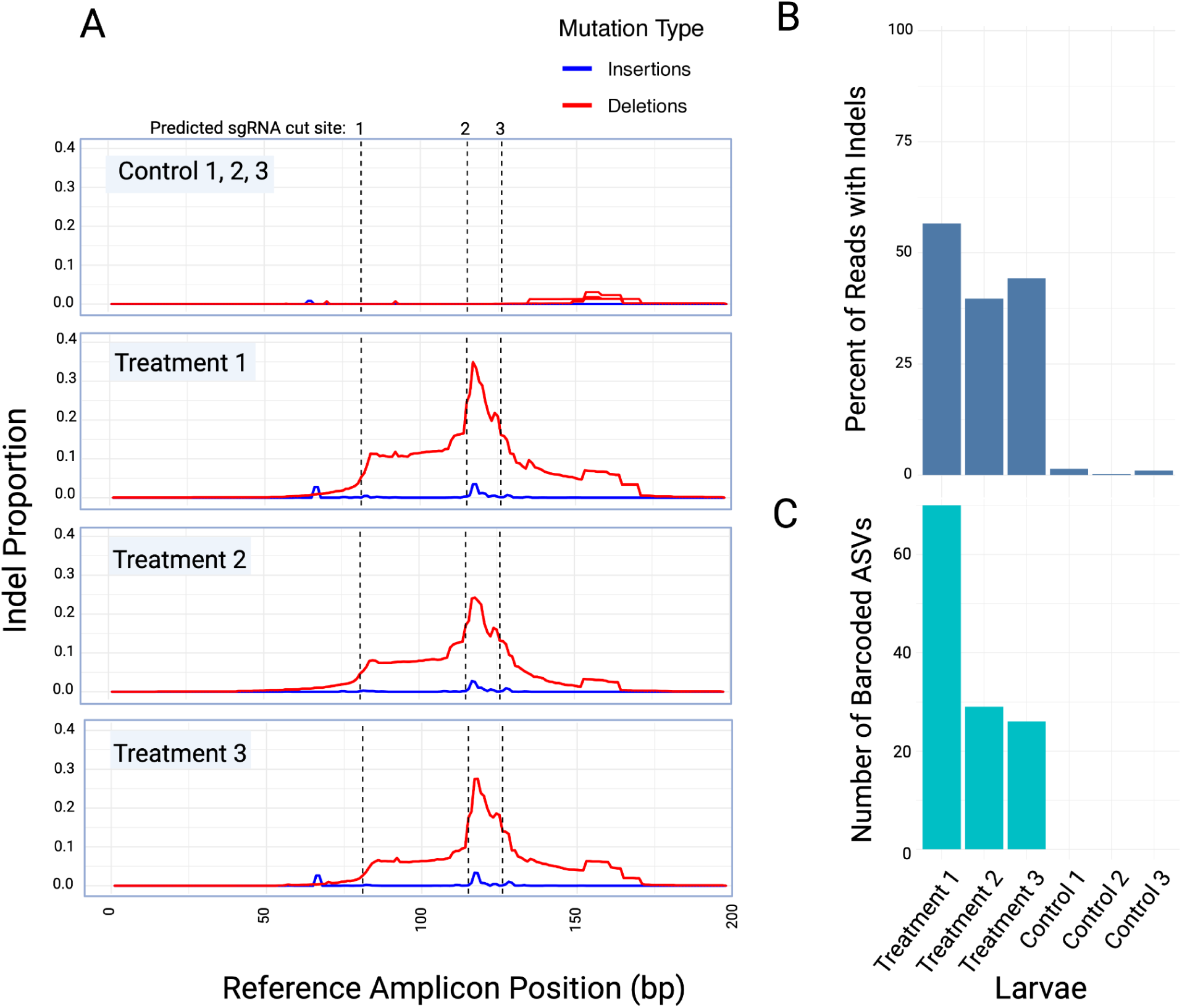
Highly effective CRISPR-Cas9 mediated genome editing of the repetitive target. (A) Proportion of insertions (blue) and deletions (red) at each bp location across the target amplicon in relation to the most dominant parental paralog. Vertical dashed bars represent predicted sgRNA cut sites. (B) Percentage of reads that contained indels for each individual relative to the most dominant WT paralog. (C) Number of ASVs in each individual that were not found in the parents.

### Widespread distribution of genetic barcodes in the *A. millepora* genome

To determine which paralogs present in the reference genome also occur in our corals, we compared amplicon sequences of the repeat from parental colonies to sequences from the reference genome. Out of the 11 reference paralogues, only three were detected in the parents. However, these three paralogs accounted for 76% (31/41) of the target sites across the genome and included the two most conserved paralogs (Fig. 3A). Sequencing results showed that we were able to successfully induce indels in all three paralogs (Fig. 3B), and that these mutated paralogs were widely distributed on nine (out of 14) chromosomes and three (out of 840) unplaced genetic scaffolds (Fig. 3C). Together, these results demonstrate wide-spread insertion of genetic barcodes throughout the genome and on multiple chromosomes.

**Fig. 3:**
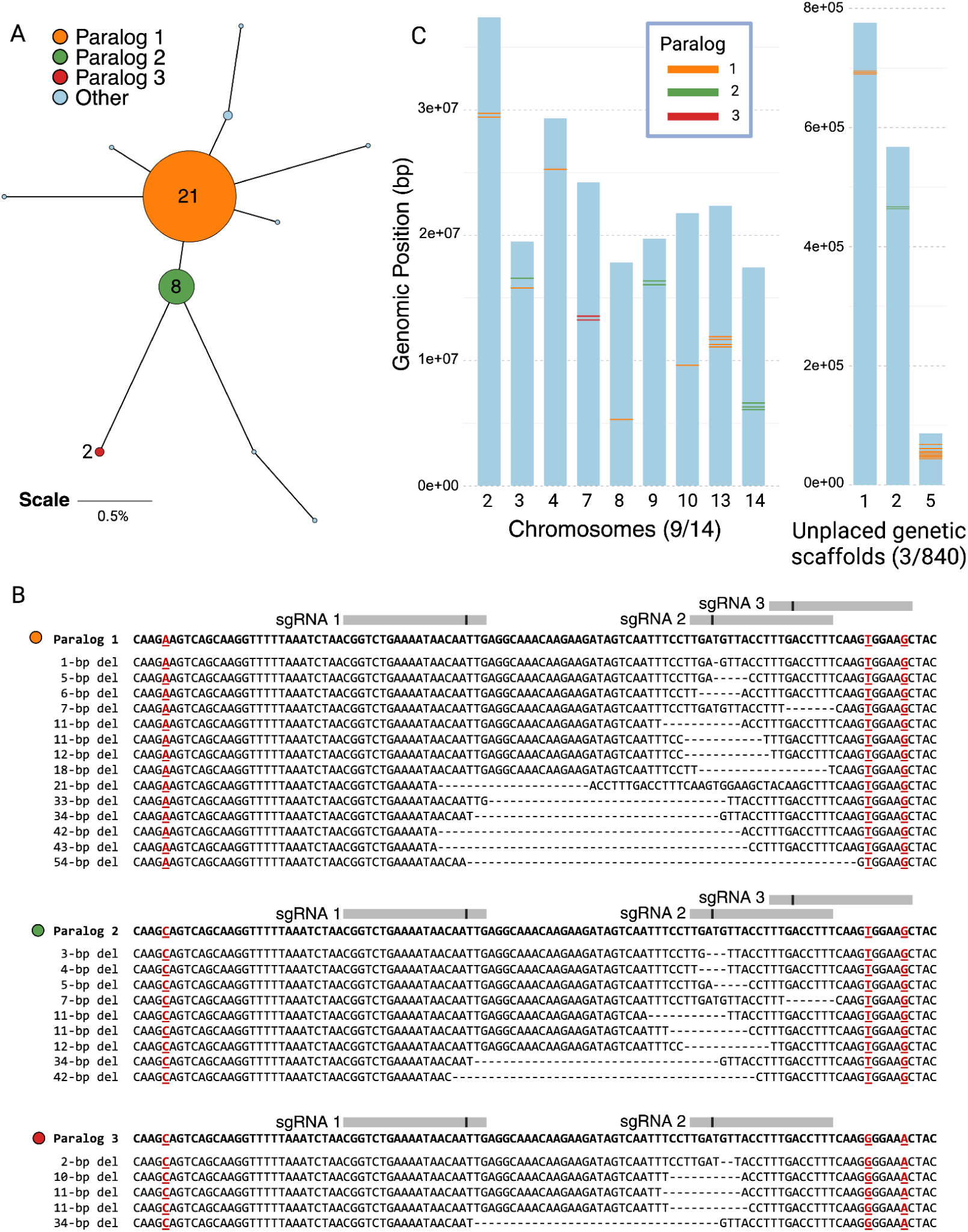
CRISPR-Cas9 genome editing of 3 distinct parental paralogs in a single individual. (A) Haplotype network showing sequence difference between 11 unique target paralogs, with 3 parental paralogs coloured. Branch lengths are proportional to percent sequence difference between paralogs. Paralog abundance is indicated for paralogs with abundance >1 and node sizes are scaled accordingly. (B) Reads from Treatment 1 larvae mapped to the three reference paralogs present in the parents. SNPs in flanking regions around indels used to assign edited sequences to a reference paralog are underlined and marked in red. Grey bars represent sgRNA binding sites with vertical black lines indicating cleavage position. Mutation type is indicated on the left of each sequence.(C) Distribution of 3 successfully mutated reference paralogs found in parents in the *A. millepora* reference genome, showing 9 (out of 14) chromosomes and 3 (out of 840) unplaced genetic scaffolds. sgRNA cut sites for respective paralogs are coloured and represented with horizontal bars.

### Multigenerational barcode inheritance

As germline cells transition from diploid to haploid in meiosis II, two scenarios can guarantee barcode inheritance: (1) if both homologous chromosomes are mutated at the same locus, all zygotes will inherit at least one edited allele; (2) if mutations are spread across many loci, the probability that at least one edited site is passed to any given gamete approaches certainty. Our most efficiently edited larvae had an average of 46% of sequencing reads containing edited sequences, leading to an estimated ∼38 unique mutations per cell (46% of 82 target loci ≈ 38). Assuming mutations are randomly distributed and conservatively disregarding linkage between target sites, the probability of guaranteeing barcode inheritance by editing at least one set of homologous chromosomes exceeds 99.99% in our most efficiently edited animals, based on combinatorial probability where L is the number of haploid target loci and *n* is the number of distinct mutations.

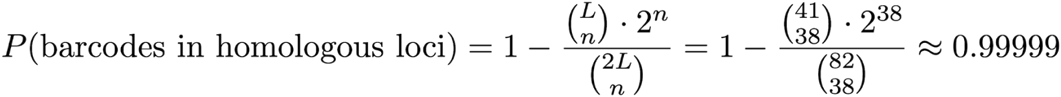

Even without editing homologous loci, we assessed whether zygotes are likely to inherit at least one barcode via recombination and chromosome segregation and whether these barcodes could persist through multiple generations. Conservatively assuming complete linkage of target sites on chromosomes/scaffolds, our repetitive region spans 14 distinct linked loci across the *A. millepora* genome. Given that each cell contains mutations in 38 of the 41 targets, zygotes would contain mutations in at least 11 of the 14 unlinked loci. We then modelled random chromosome segregation during meiosis II over multiple generations to estimate the percentage of offspring in each generation that would inherit at least one barcoded locus. The probability of inheriting at least one mutated locus is given by,

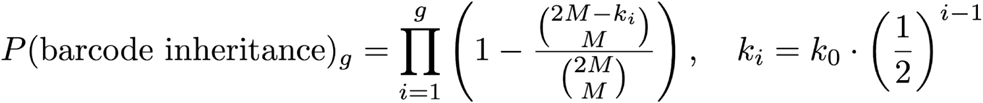

which assumes each zygote inherits one haploid set of unlinked target loci (*M*) and that the number of barcoded loci (*k*) halves each generation (*g*) (following mendelian inheritance). Thus, with 11 of 14 loci mutated (*k_0_* = 11, *M* = 14), the proportion of offspring that inherit barcodes is greater than 99% in the first two generations and remains high over multiple generations (Table 1). Taken together, these analyses suggest that barcode transmission to offspring is highly probable and can persist over multiple generations in our most efficiently edited larvae.

**Table 1.**
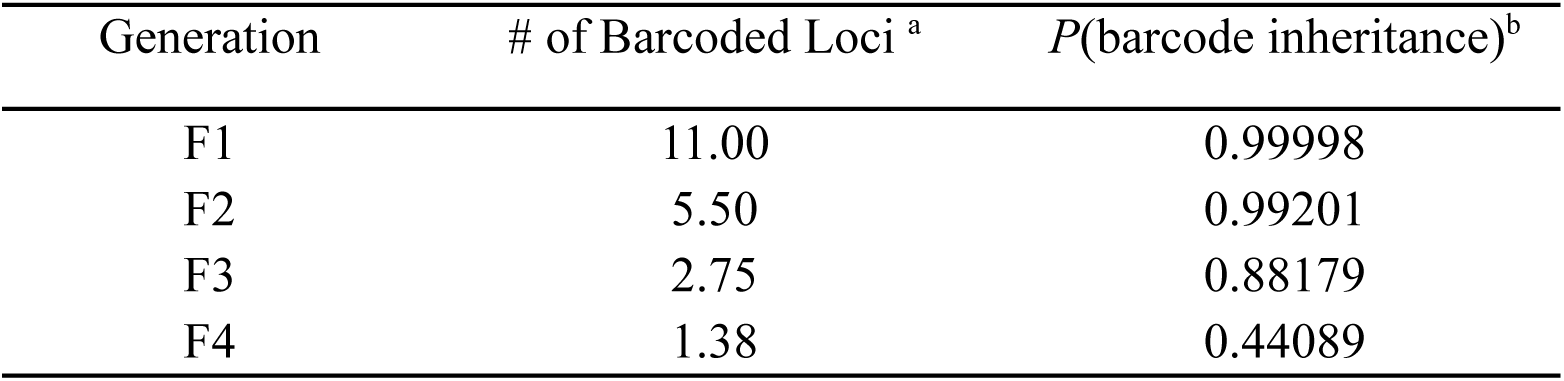
Estimated proportion of offspring that inherit barcodes over multiple generations. ^a^ Average number of mutated loci in each parental generation (*k_i-1_*) assuming that *k*_0_ = 11. ^b^ Approximate proportion of offspring in each generation that will inherit at least one unique barcode, where 2M - *k_i_*was rounded to the nearest whole integer.

| Generation | # of Barcoded Loci <sup>a</sup> | $P(\text{barcode inheritance})^b$ |
| --- | --- | --- |
| F1 | 11.00 | 0.99998 |
| F2 | 5.50 | 0.99201 |
| F3 | 2.75 | 0.88179 |
| F4 | 1.38 | 0.44089 |

### Detecting genetic barcodes with eDNA

Having successfully induced genetic barcodes in *A. millepora* corals, we examined whether barcodes could be tracked using environmental DNA (eDNA). To do this, six new treatment larvae were kept in 300ml of seawater for 24 hrs, after which the larvae were removed and the remaining seawater was filtered. Identical samples were taken for control injected larvae. Amplicon-sequencing of the repetitive site from seawater DNA showed that water from treatment larvae carried DNA with indels at the sgRNA target site (Fig. 4). We analysed the proportion of insertions and deletions across the editing window for the 6 injected larvae, which revealed the larvae exhibiting a range of indel frequencies (Fig. 4A). As we would expect the indel frequencies found in the true eDNA sample to be the approximate average of indel frequencies from the six treatment larvae, we created a simulated eDNA sample by randomly sampling and concatenating reads from the 6 injected larvae. Analysis revealed the indel pattern from the simulated eDNA sample closely mirrors that of the real eDNA sample, with the magnitude of indel proportions across the editing window being nearly identical for real and simulated samples (Fig. 4B). Furthermore, we analysed the percent of barcoded reads between the eDNA sample and the injected larvae and similarly found that the percent of barcoded reads in the eDNA was an approximate average of the percent of barcoded reads between larvae (Fig. 4C).

**Fig. 4:**
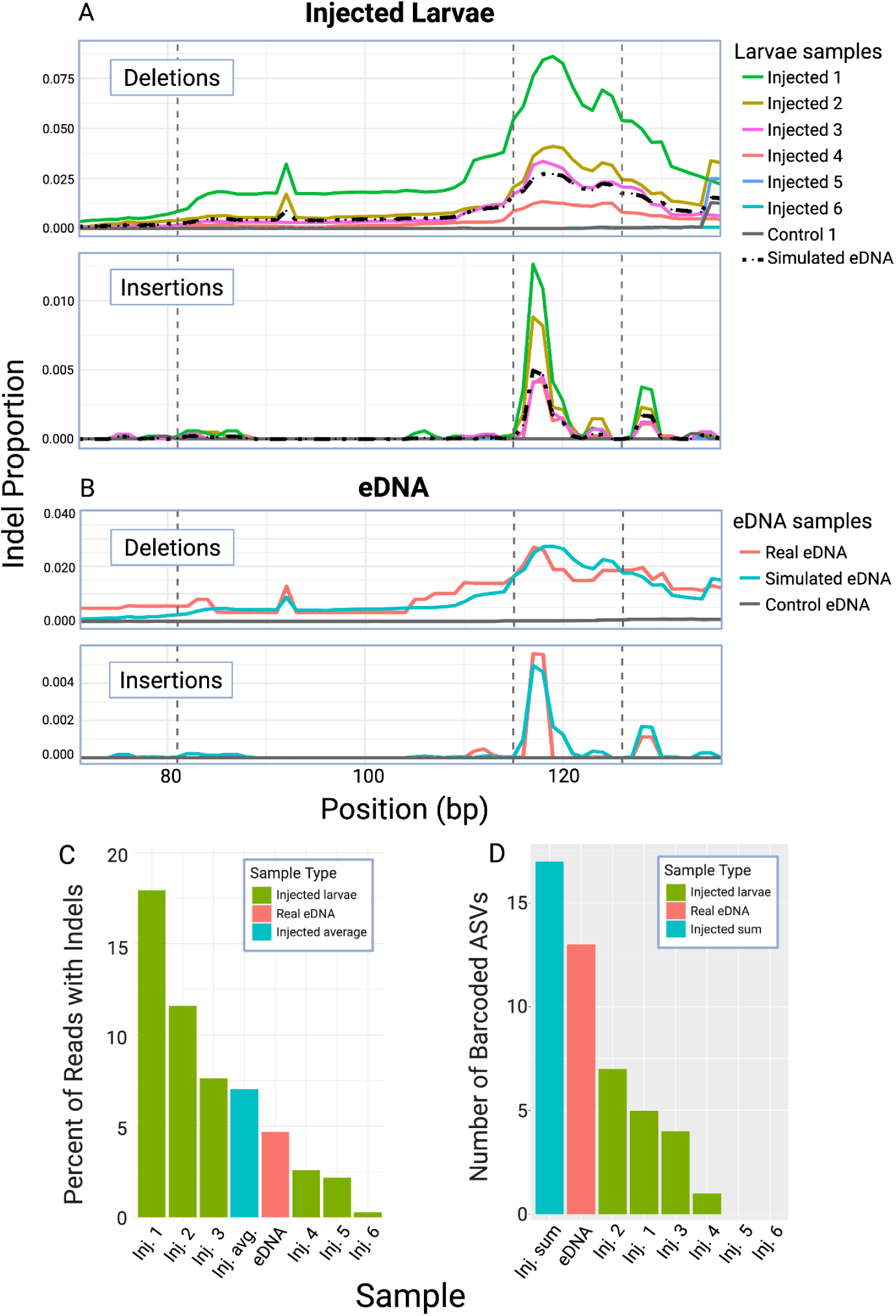
Detection and quantification of barcoded sequences in eDNA. (A & B) Proportion of insertions and deletions at each bp location in the sgRNA target window. Vertical dashed bars represent predicted sgRNA cut sites. (A) Indel proportions of injected larvae and their represented average (simulated eDNA) compared to control. (B) Indel proportions of the real eDNA and simulated eDNA samples compared to control. (C) Percentage of filtered reads that contained indels from injected larvae, real eDNA, and injected average. (D) Number of barcoded ASVs found in each sample compared between injected larvae, real eDNA, and injected sum.

While the proportion of barcoded sequences in the eDNA sample was close to the average of the larvae samples, we would expect the number of distinct barcode sequences in the eDNA sample to be a sum of the larvae samples, given the eDNA sample was able to efficiently trace DNA shed from each larva. To explore this, we compared the quantity of unique barcodes between the eDNA sample and the contributing larvae. As expected, we found that the eDNA sample had substantially more unique ASVs than each of the 6 donor larvae, with the eDNA sample containing 13 barcoded ASVs and the larvae having a combined total of 17 barcoded ASVs (Fig. 4D). Together, these results not only demonstrate that we can detect our genetic barcodes from eDNA, but also that these barcodes follow predictable patterns that reflect their source population.

## Discussion

In response to the global species loss, there has been a widespread expansion of efforts to manage and restore affected populations^4–9^. However, existing techniques for monitoring species used in population restoration remain limited in their ability to track restored individuals and their offspring across broad spatial and temporal scales^13,14,23^. Here, we demonstrate that PCR-detection of CRISPR-Cas9 induced genomic barcodes in environmental DNA (eDNA) offers a novel framework to address these limitations. We provide a system that creates a genomic tag in corals that allows for tracking of individuals and their offspring non-invasively using eDNA, providing a permanent, heritable, and potentially scalable approach to monitoring.

In controlled aquaria, barcode sequences recovered from eDNA showed indel frequencies and patterns that closely matched those of the edited individuals contributing to the sample (Fig. 4). eDNA-based detection provides a non-invasive method for genetically characterizing the presence of treated coral without the need for sacrificial sampling. This is particularly important for genetic screening in early life stages, as non-lethal tissue-based sampling typically requires larvae to be reared to later developmental stages. Furthermore, as eDNA monitoring can be a cost-effective, scalable method for population surveillance^47–50^, detecting permanent genetic barcodes through seawater sampling could provide a powerful tool for monitoring survival and spread of restored corals at scale. However, future experiments are needed to validate the feasibility of detecting these barcodes in field-collected eDNA, where detection can be confounded by environmental noise, DNA degradation, and dilution effects^51–54^. Furthermore, while eDNA methods show promise to allow scalable monitoring, current CRISPR-Cas9 transfection techniques (such as microinjection) used in corals are limited in scale (∼1,000 zygotes can be injected by a single person per night of spawning). Developing other transfection methods in corals, such as chemical transfection or electroporation, which are developed in other species, would greatly increase the scalability of this approach.

Our analyses indicate that barcode transmission to offspring is highly likely and can persist for multiple generations in the most efficiently edited larvae. This provides a powerful tool to address questions of larval dispersal and reef connectivity. Although individuals with lower editing efficiency may transmit barcodes to fewer descendants, inheritance is still probable in a subset. However, these theoretical predictions need to be tested empirically through the maturation and spawning of edited corals to confirm transgenerational barcode inheritance.

While targeting repetitive loci offers advantages, such as increased editing efficiency and improved inheritance probability, it also introduces potential complications as it also dilutes the copy frequency of any single barcode. For instance, each barcode introduced at the single-cell stage in a single-copy locus would represent 50% of amplicon sequences from an individual, whereas each unique barcode introduced in a 41-locus target would represent only ∼1.2% of amplicon sequences. This effect becomes more pronounced in mixed population eDNA samples. For example, if one barcoded larva is pooled with nine WT larvae, the relative read frequency of each unique barcode sequence drops further - from ∼1.2% to ∼0.12%. This problem would only increase as reef populations become more diverse and future research should explore strategies to mitigate this issue. One promising approach to do so is in vitro CRISPR-based depletion, in which eDNA samples are treated with sgRNA/Cas9 targeting wild-type sequences^55–57^. This would selectively digest non-barcoded alleles, effectively enriching barcode sequences in downstream sequencing libraries, improving detection sensitivity.

In conclusion, our study demonstrates that CRISPR-Cas9 genome editing and eDNA monitoring has the potential to be integrated into a practical framework for tracking corals and their offspring at scale. This capability has the potential to allow unprecedented insight into the persistence, dispersal, and reproductive output of restored individuals, and could be applied to restoration practices in a wide range of taxa. As climate change intensifies and species loss accelerates, these kinds of precision-tracking tools will be critical for evaluating which interventions are working, how well and where. At the same time, deploying genetic technologies in natural systems demands scrutiny. As these technologies become more powerful, it will be essential to pair their application with robust empirical validation and a measured, ecosystem-based approach to conservation.

## Methodology

### Animal collection, spawning and larvae husbandry

Individual *A. millepora* colonies were collected in mid-November 2024 from the Palms Islands on the Great Barrier Reef (Permit G21/38062.1) and brought to holding tanks at the Australian Institute of Marine Science National Sea Simulator. Corals were kept in flow through seawater tanks during the day and isolated in individual tanks at night. The corals spawned on November 21^st^ at around 21:00pm. Egg and sperm bundles were collected from each colony and washed with 125 µm filters. Gametes from each colony were kept separate to allow staggered fertilizations, which were made by combining 10 mL of sperm at ∼10^6^ cells/mL with ∼300 pooled eggs in 300ml of seawater and incubating for 15 min at 27°C. Staggered fertilizations were made every 1 hour for 6 hours post spawn, which allowed zygotes to be transfected at the single cell stage, as the first cell cleavage occurs 30-90 minutes post fertilization.

### Identifying a repetitive target

A repetitive, non-coding DNA sequence was identified in the *Acropora millepora* genome using *RepeatMasker*^58^. A BLASTn was then performed querying the repeat sequence against the *A. millepora* reference genome on NCBI (Accession PRJNA633778, Ref-seq GCF_013753865.1). This BLASTn yielded many hits, none of which overlapped with any known genes or other annotations. PCR primers were then designed to amplify a 234 bp region spanning the repeat, and validated to ensure the repetitive sequence could be successfully amplified.

### Design and synthesis of sgRNAs

CRISPR-Cas9 guide RNAs were designed to target the repetitive sequence using the online guide design tool ChopChop^59^. sgRNA that had the highest editing efficiency across sgRNA design tools were chosen and ordered from IDT’s gBlocks gene fragments service in the form of a DNA template containing a T7 promoter, a target sgRNA sequence, a Cas9 scaffold sequence, and a poly T tail (Fig. S1). gBlock templates were reconstituted in nuclease free water to a concentration of 300 nM, and 3 μL were used in a 30 μL in vitro transcription reaction using a HiScribe T7 RNA Synthesis Kit (NEB no. E2050S) to synthesize sgRNAs. Synthesized sgRNAs were purified using an RNA clean and concentrator kit (Zymo Research no. R1013) and their quality and size was verified using gel electrophoresis and quantified using a NanoDrop2000 Spectrophotometer (ThermoFisher). sgRNA/Cas9 ribonucleoprotein complexes were then generated by incubating each sgRNA with Cas9 in separate reactions as described previously^44^.

### Chromosome mapping and paralog identification

The genomic locations of the target repeat in the *A. millepora* genome were identified using a multistep BLASTn. First, a BLASTn querying the forward PCR Primer against the reference genome was performed. Next, flaking regions around each forward primer hit were extracted and a BLASTn querying the reverse PCR primer was performed on these flanking regions. This yielded a total of 41 target regions in the *A. millepora* genome that should be amplified by the PCR primer set. The sequences contained within the primer set at the 41 target sites were then extracted and clustered with 100% sequence identity using *CD-HIT-EST* to identify 11 unique paralogs present in the reference genome. Sequence similarity was then visualized in a haplotype network using a distance matrix generated with Tree Builder from Geneious Prime. Next, a BLASTn was performed querying the 3 sgRNAs against 11 reference paralogs, which showed that all 11 reference paralogs making up the 41 target regions had a 100% sequence match for at least 2 of the 3 sgRNAs. These 41 target sites were then mapped to the reference genome to quantify the distribution of the repetitive target across chromosomes.

### Microinjection and identification of successfully injected larvae

Transfection of zygotes with CRISPR-Cas9 RNP was performed via microinjection as described previously^44,45^. In short, CRISPR-Cas9 RNP was injected along with a fluorescent dextran indicator (Alexa Fluor 488-labeled dextran; Invitrogen no. D22910). WT larvae were also injected but with an injection mix that lacked any sgRNA. Coral zygotes were injected to a volume of approximately 10-20% of the total volume of the zygote under a fluorescent microscope (GFP filter, excitation 488 nm). Injected larvae were sorted for successful injection 12 hours post-fertilization by comparing fluorescence of injected larvae to WT controls. Larvae that were fluorescent were considered successfully injected, while non-fluorescent larvae were classified as unsuccessfully injected.

### Sample fixation, eDNA collection, and sequencing

Successfully injected larvae were kept in 300 ml tubs of autoclaved seawater at 27 °C until they were 3 days old. Larvae used for the initial gene editing analyses were fixed at 3 days old by removing them individually from the seawater tubs and storing them in 100% ethanol at −20 °C. To produce both treatment and control eDNA samples, 6 successfully injected larvae of 3 days old were transferred together to a new 300 ml tub of autoclaved seawater and kept for 24 hrs. The larvae were then individually removed from the shared tub and stored in 100% ethanol at −20 °C. The leftover seawater (∼300 mL) was then filtered through a 0.22 μm filter to collect eDNA shed by the 6 larvae. Tissue samples were also taken from all parent colonies by sampling a nubbin of ∼1 cm^2^ from each colony and storing them in 100% ethanol at room temperature. DNA was extracted from all samples using the Maxwell RSC PureFood GMO and Authentication Kit (Promega no. AS1600). PCR was performed to amplify CRISPR-Cas9 target regions from the DNA extracts of all samples using PCR Primers with Illumina adaptors and the following parameters: one cycle of 95 °C for 10 min; 30 cycles of 95 °C for 30 s, 55 °C for 30 s, and 72 °C for 30 s; and one cycle of 72 °C for 10 min. The resulting amplicons were then verified with gel electrophoresis and sequenced at AGRF in Brisbane using next-generation amplicon sequencing with the Illumina MiSeq system (150bp Paired End reads). As the eDNA sample contained a mix of DNA from six larvae, the eDNA sample was sequenced at a higher depth than tissue samples to normalize the effort used to detect any given sequence per individual, with the eDNA sample having 25,109 reads and tissue samples having a mean ± SD of 5,844 ± 1,325 reads.

### Read processing and mutation quantification

Paired end MiSeq reads were first trimmed for quality using *cutadapt* (Phred >35) for all samples. The simulated eDNA sample was generated by randomly sampling reads from the 6 injected larvae matching the real eDNA sample and concatenating them into a new file. 4,185 reads were sampled from each of the 6 larvae so that the read count of the resulting simulated eDNA sample matched the read count of the real eDNA sample. CRISPR-Cas9 mediated insertions and deletions (indels) were identified in all samples using CRISPResso, which merges paired-end reads, aligns them to a reference amplicon sequence, and quantifies the numbers of reads with indels near the predicted Cas9 cut site, generating outputs that quantify indel frequencies, sizes, and positions throughout the amplicon for each individual. CRISPResso was run in batch with parameters to ignore substitutions, as they can less confidently be ascribed to a CRISPR-Cas9 mediated cut than indels. Furthermore, possible indels that were supported by less than 5 reads were discarded. To quantify the number of distinct mutations generated in each individual, filtered MiSeq reads from all samples were clustered into Amplicon Sequence Variants (ASV) using the Dada2 pipeline^60^ and further filtered using LULU^61^ with 98% sequence identity. An ASV that was present in a larvae sample but not present in Parent samples was considered a unique ASV.

### Mapping barcodes to paralogs and chromosomes

To identify the paralogs shared between the reference genome and our corals, a BLASTn was performed querying filtered reads from parent samples against the 11 reference paralogs using 100% identity. Reference paralogs that had hits from > 2% of reads from a single parent were considered present, which was the case for 3 of the 11 reference paralogs. Filtered MiSeq reads from Treatment 1 were then clustered with 100% identity using *CD-HIT-EST* and mapped to the three reference paralogs, as SNPs in flanking regions around CRISPR induced indels could assign edited sequences to a reference paralog. Clusters containing less than 10 reads were removed and only sequences that perfectly mapped outside of the CRISPR induced mutation, contained a noncomplex mutation (only a single deletion or insertion), and could uniquely map to only one paralog were included.

## Acknowledgments

We thank A. Sedgwick, E. Pfeffer, H. Razali, and G. Luther for their contributions during coral spawning and E. Meier for her guidance during microinjection. Funding for this study was provided by an internal grant from the Australian Institute of Marine Science (to L.B. and L.T.). Coral collection was conducted under Great Barrier Reef Marine Park Authority (GBRMPA) permit G21/38062.1.

## Author contributions

L.B., P.C., L.T., and M.M. collaboratively designed the study. M.M. and V.M. conducted the experiments. L.T., Y.S. and R.L. provided laboratory space, equipment, and logistical support. M.M. performed the formal analysis and generated the visualizations. M.M. wrote the original draft and all authors contributed to reviewing and editing the manuscript.

## Competing interests

The authors declare no competing interests.

## Data availability

New sequencing data for amplicons of the repeat region have been uploaded to the NCBI Sequence Read Archive (SRA) at accession number PRJNA1290875. The *A. millepora* reference genome used in this study can be found on NCBI at accession number PRJNA633778.

## Supplemental Figures

**Fig. S1:**
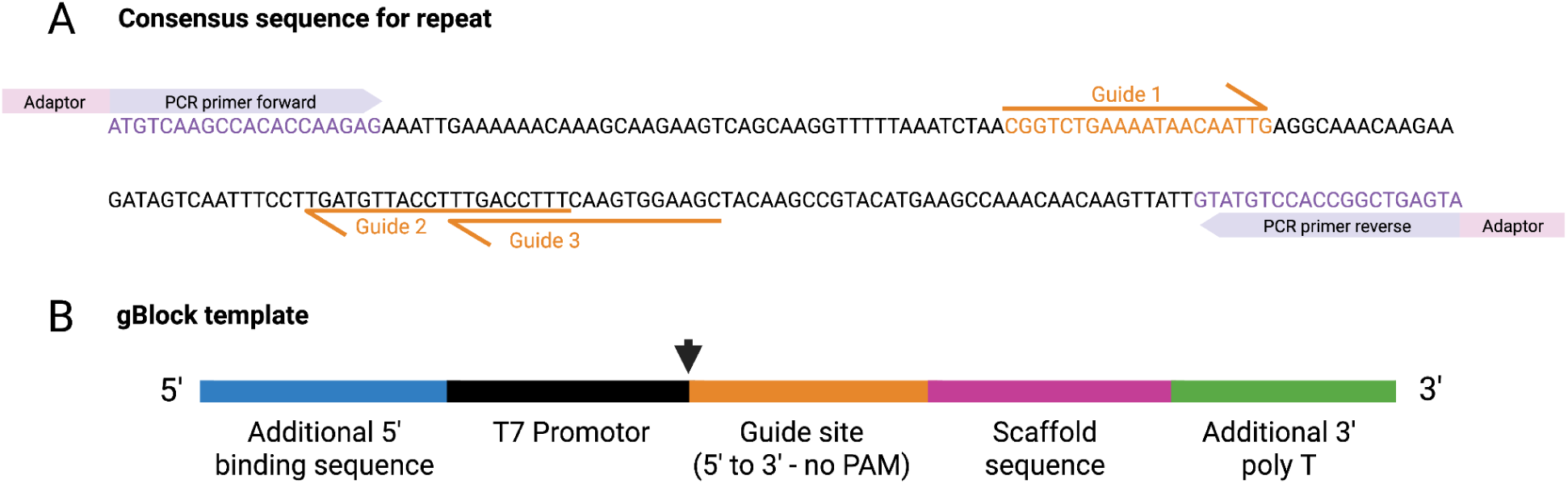
Single guide RNAs and their gBlock templates. (A) The most abundant variant of the repetitive sequence and locations of its corresponding PCR primers + Illumina adaptors (purple + pink) and sgRNA target sites (orange). (B) Template format of gBlocks gene fragments for complete sgRNA with 5’ binding sequence (blue), T7 promoter (black), sgRNA guide site (orange), scaffold sequence for Cas9 binding (pink), and a poly T tail (green). Arrow indicates transcription start site.

**Figure S2.**
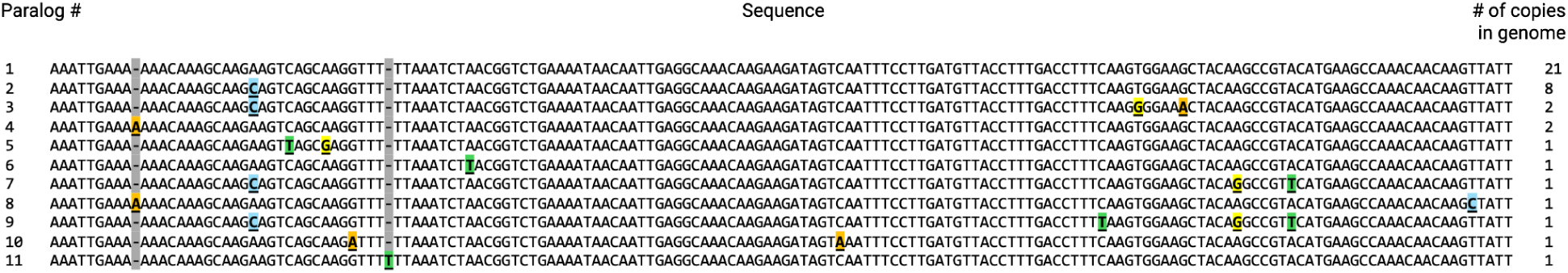
Sequence alignment and genomic copy number of 11 reference paralogs. Reference paralog number, paralog sequence, and genomic copy number are shown from left to right for each paralog. SNPs in each paralog that differ from Paralog 1 are underlined and base pair changes are coloured respectively (A-orange, G-yellow, C-blue, T-green). Insertions are tracked with a dash and highlighted in grey.

**Fig. S3:**
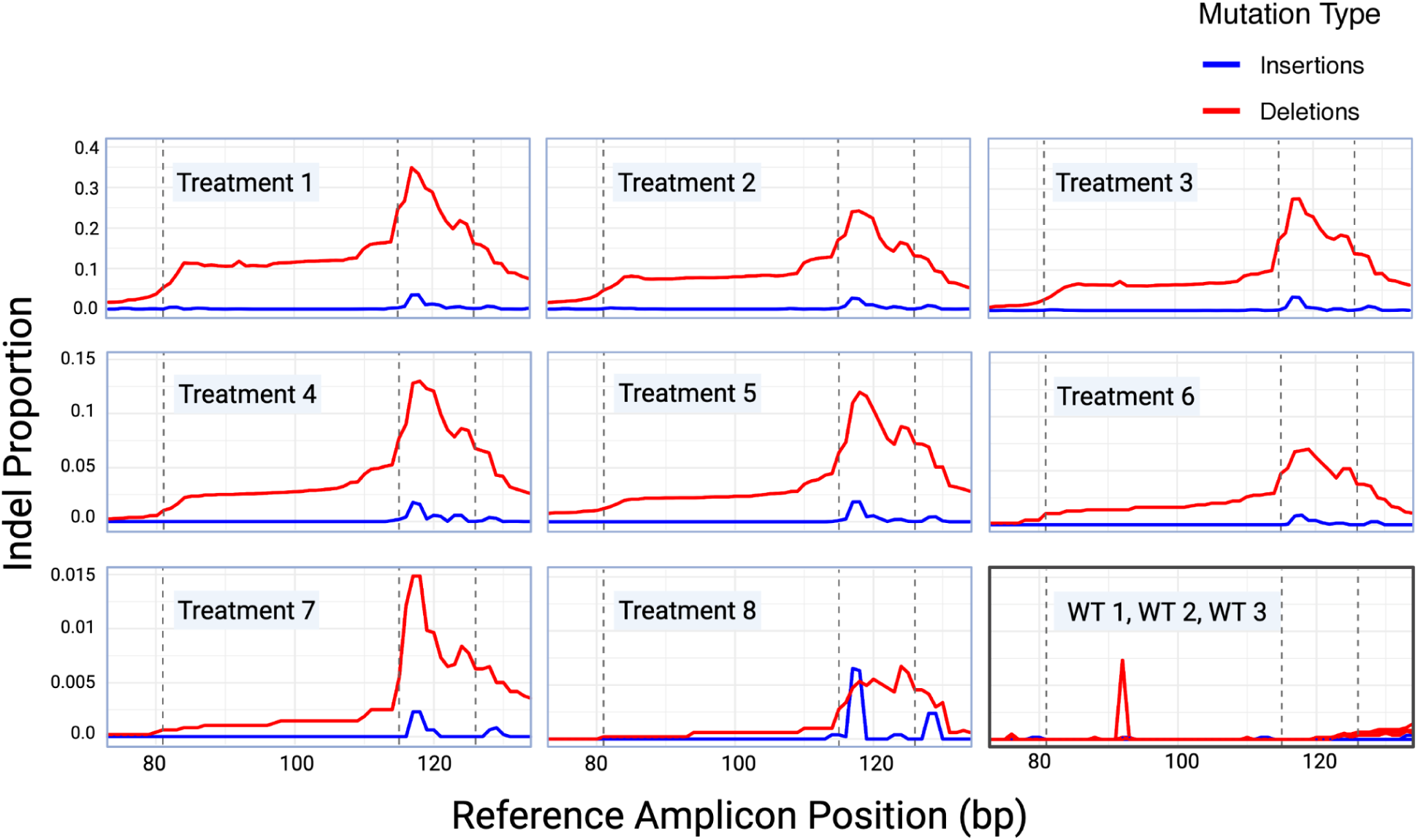
CRISPR-Cas9 mediated genome editing of our *A. millepora* repetitive target in 8 injected larvae. Plots show the proportion of insertions (blue) and deletions (red) at each bp location across the target amplicon. Vertical dashed bars represent predicted sgRNA cut sites. Plots for the 8 treatment larvae are labelled and boxed in blue. Indel proportions for the three WT larvae are shown together in one plot boxed in black (bottom right). Note the different Indel proportion scale for each row.

**Table S1:**
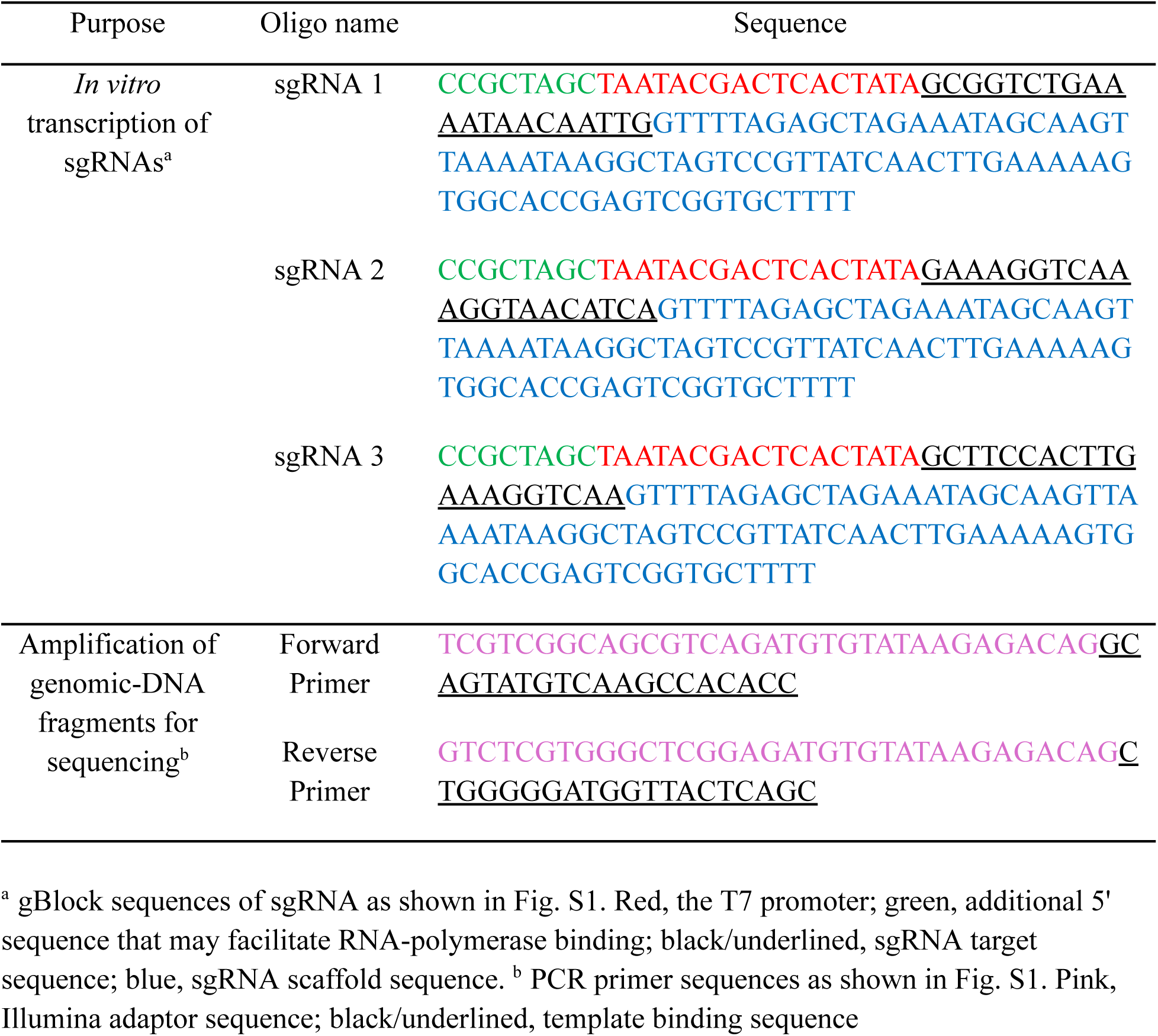
Oligonucleotides used in this study.

## Notes

### Competing Interest Statement

The authors have declared no competing interest.

